# Single-cell metabolic profiling reveals subgroups of primary human hepatocytes showing heterogeneous responses to drug challenge

**DOI:** 10.1101/2022.06.08.495252

**Authors:** E. Sanchez-Quant, M. L. Richter, M. Colomé-Tatché, C.P. Martinez-Jimenez

## Abstract

Xenobiotics are primarily metabolized by hepatocytes in the liver, and primary human hepatocytes (PHHs) are the gold standard model for the assessment of drug efficacy, safety and toxicity in the early phases of drug development. Recent advances in single-cell genomics have shown liver zonation and ploidy as main drivers of cellular heterogeneity. However, little is known about the impact of hepatocyte specialization on liver function upon metabolic challenge, including hepatic metabolism, detoxification, and protein synthesis. Here, we investigate the metabolic capacity of individual human hepatocytes *in vitro*, and assess how chronic accumulation of lipids enhances cellular heterogeneity and impairs the metabolisms of drugs. A phenotyping five-probe cocktail was used to identify four functional subgroups of hepatocytes that respond differently to drug challenge and fatty acid accumulation. These four subgroups display differential gene expression profiles upon cocktail treatment and xenobiotic metabolism-related specialization. Notably, intracellular fat accumulation leads to increased transcriptional variability and diminished the drug-related metabolic capacity of hepatocytes. Our results demonstrate that, upon a metabolic challenge such as exposure to drugs or intracellular fat accumulation, hepatocyte subgroups lead to different and heterogeneous transcriptional responses.

## INTRODUCTION

Single-cell sequencing technologies have revealed a wealth of information regarding cellular heterogeneity in health and disease in multiple tissues [1–8]. In particular, single-cell RNA sequencing enables the identification of groups of cells in a population with similar transcriptomic profiles, which is generally associated with similar functionality [9, 10]. In the liver, single-cell transcriptomic analyses have shown novel cell types and states involved in the development and progression of liver disease [7, 11], as well as in the transcriptomic responses to xenobiotics [11]. However, drug toxicity and safety are generally assessed in primary human hepatocytes (PHH) as the gold standard model to study the drug metabolism in humans, which are widely considered a seemingly homogeneous population of cells [12, 13]. Whether all hepatocytes share the same functional molecular phenotype *in vitro* remains unknown.

In the liver, drug and xenobiotic metabolism is mainly performed by hepatocytes, the major and predominant cell type of the parenchyma [14–16]. The hepatic metabolism of drugs occurs in three phases. In phase I, oxidation, hydrolysis, reduction and cyclization reactions are catabolized mainly by the cytochrome P450 (CYP450) superfamily of monooxygenase enzymes. The main members are the isoforms CYP1A2, 2C9, 2C19, 2D6 and 3A4, that account for the metabolism of around 70-80% of the clinically available drugs [17–19]. The expression of these cytochromes is induced by the presence of their substrate compounds, which are indirectly used as a measure of liver metabolic capacity [10, 20]. These substrates act as hepatocyte probes when administered to phenotype and monitor the cytochrome enzymatic activity, formally known as the “phenotyping cocktail approach” [21–27]. Phase II comprises conjugation reactions catabolized mainly by transferase enzymes and are purposed to hydrophilize compounds for their elimination [28]. During phase III of xenobiotic metabolism, conjugated compounds are excreted out of the cell through active transmembrane transporters [29]. The cytochrome superfamily consists of nearly 60 members in humans (Human Genome Project 2013), which may be expressed differently in individual cells when challenged by xenobiotics, leading to cellular heterogeneity that remains concealed in a bulk analysis.

An additional potential source of cellular heterogeneity is the intracellular accumulation of triglycerides in hepatocytes, known as hepatic steatosis [30, 31]. This is a hallmark of non-alcoholic fatty liver disease (NAFLD) and is generally associated with metabolic dysfunction, inflammation and risk of fibrosis [32, 33] [34, 35]. In culture, intracytoplasmic fat accumulation can be modeled by incubating primary human hepatocytes with free fatty acids (FFA) to mimic benign chronic steatosis [36–38]. However, lipid accumulation is heterogenous in regards of number and size of lipid droplets [39] and it still remains unclear whether all cells respond to lipid accumulation in a coordinated manner.

Moreover, the co-administration of five or more drugs (polypharmacy) is associated with a higher incidence of adverse drug reactions (ADR) and drug-induced liver injury (DILI) [40] [41–44]. Importantly, a higher incidence of DILI has been reported in patients suffering from NAFLD [45–47]. Therefore, the precise molecular pathways commonly dysregulated between chronic accumulation of lipids and drugs metabolism remain unexplored at cellular resolution.

Here, we first dissect concealed cellular heterogeneity of primary human hepatocytes using scRNA-seq upon metabolic and drug challenge, and identify four distinct subgroups of hepatocytes which are present *in vitro* and *in vivo*. We further characterize their metabolic profile in response to a five-drug cocktail, highlighting their differential xenobiotic metabolism. Additionally, we mimic hepatic steatosis *in vitro* and discover that intracellular accumulation of lipids leads to increased loss of expression and transcriptional variability. Taken together, we show that lipid accumulation in hepatocytes leads to subgroup-specific impairment of multiple metabolic pathways. Our results illustrate that changes in cellular heterogeneity and intracellular lipid accumulation increase transcriptional noise and susceptibility to adverse drug reactions in humans.

## RESULTS

### Single-cell RNA-seq reveals four major subgroups of hepatocytes showing cellular heterogeneity and functional specialization in primary human hepatocytes

Liver function is compartmentalized along the liver lobule, and the maintenance of liver function is regulated by liver-enriched transcription factors [1, 4, 48–51]. Primary human hepatocytes are considered the gold standard model to predict drug responses in humans, but they are characterized by phenotypic instability in culture and rapid loss of the hepatic phenotype [12, 13, 52–55]. Therefore, we aimed to investigate the level of cellular heterogeneity that remains in a seemingly homogeneous population of primary human hepatocytes (PHHs), and how this heterogeneity might affect pharmaco-toxicological studies [56–58].

Cryopreserved PHHs from four donors were used as *in vitro* model to characterize the metabolic profile of individual hepatocytes in response to a drug challenge and chronic accumulation of fat (Figure 1A). The hepatocytes were plated for 6 hours and incubated for 66 hours with either vehicle (DMSO) or a five-drug cocktail. Chronic accumulation of fat was achieved by incubation with free fatty acids (FFA) corresponding to a 200 µM mixture of a 2:1 ratio of oleic acid to palmitic acid for 72 hours. In brief, four different conditions were studied: i) Vehicle (DMSO 0.5% v/v); ii) Cocktail (66h five-drug cocktail incubation); iii) FFA (2:1 oleic:palmitic acid), and iv) FFA+Cocktail (combination of FFA incubation and five-drug cocktail) (Figure 1A). Donors were healthy males between 18 and 57 years, with a Body Mass Index (BMI) classified as normal, non-diabetic, and representing the most common age range commercially available (Supp. Table 1). After isolation of viable cells by a quick and non-harsh dead cell removal step, single-cell RNA-seq was immediately performed using 10x Genomics (Methods).

**Figure 1:**
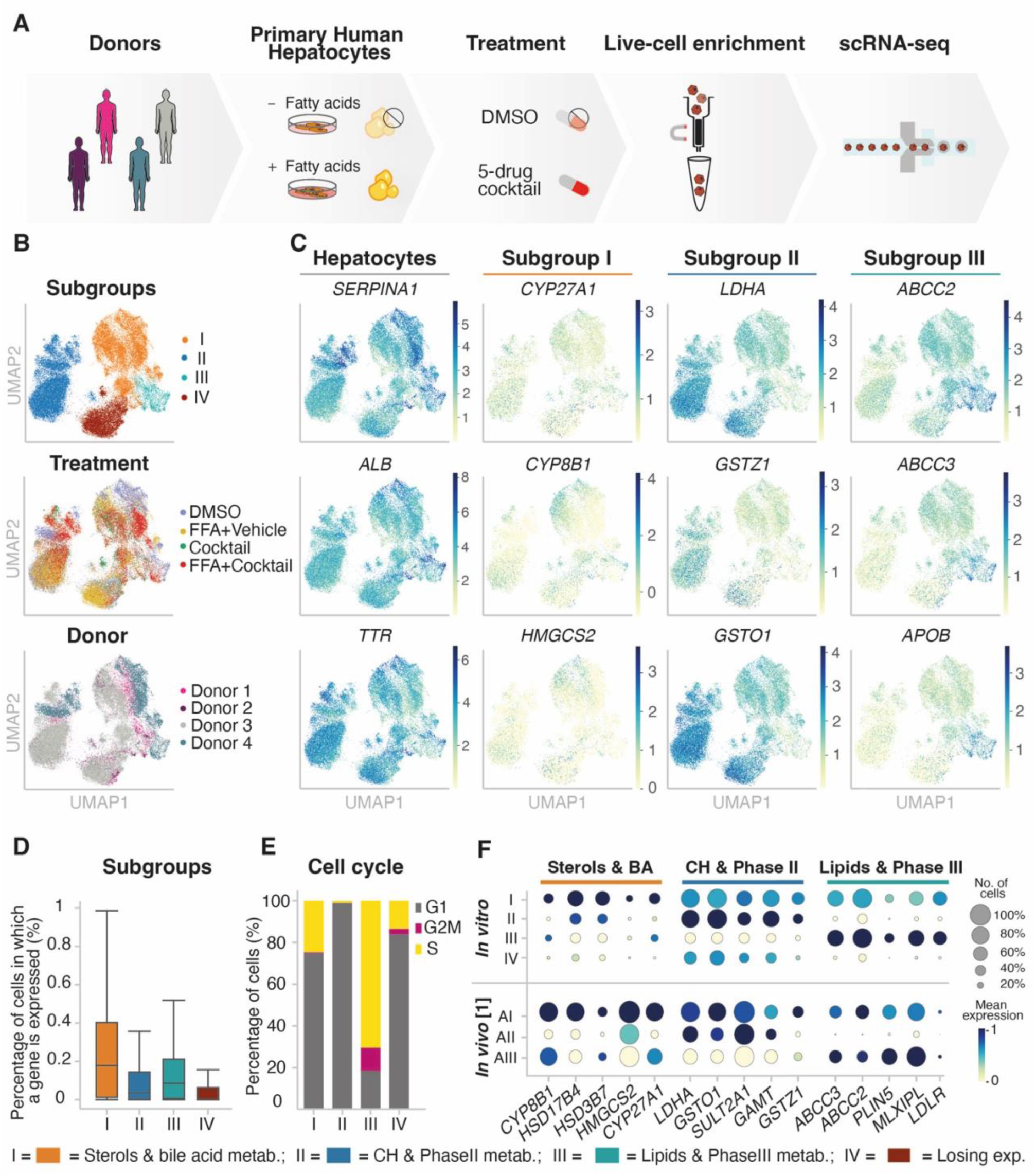
Transcriptomic profiling reveals four subgroups of PHHs independently of donor and treatment condition. **(A)** Overview of experimental design. Purified cryopreserved PHHs from four human donors were plated, incubated with or without free fatty acids to model hepatic fat accumulation and with or without a phenotypic 5-drug cocktail (Sanofi-Aventis). **(B)** UMAP colored by subgroup, treatment, and donor showing that the annotated subgroups are found throughout all donors and conditions. **(C)** UMAP colored by expression levels of - from left to right: i) Key hepatocyte marker genes; ii) Bile acid and sterol metabolism; iii) Carbohydrate and phase II metabolism; iv) Lipid and phase III metabolism marker genes in four subgroups of hepatocytes. **(D)** Boxplot showing the percentage of cells in which a given gene is expressed, colored by the identified subgroups. **(E)** Bar plot showing the percentage of cells per subgroup assigned to phases G1, S, and G2M by performing cell cycle analysis using *cyclone.* **(F)**Dot plot highlighting marker gene expression in four subgroups of hepatocytes identified *in vitro* (top) and *in vivo* (Aizarani *et al*. 2019). Subgroup marker gene expression was grouped by aggregated *Louvain* clusters (dot size: fraction of cells in group; color scale: mean expression in group).

A total of 38,232 high-quality hepatocytes with an average of 2,550 genes per cell were analyzed. Lenient doublet filtering was performed to account for polyploid hepatocytes (Methods). The variation in the number of cells profiled per donor was attributed in part to sample viability, technical differences in cell capture rates in each scRNA-seq run, and sequencing depth (Supp. Figure 1, Methods).

Across all four donors, *Louvain* clustering led to the detection of heterogeneity separating four subgroups of hepatocytes, independently of the treatment (Figure 1B, Supp. Figure 1B and C, Methods). *Harmony* [154] was used to correct for batch effects, and *Louvain* clustering was performed on the combined data set (Supp. Figure 2A). Overall, hepatocyte purity was confirmed by the expression of the hepatocyte-specific genes *ALB*, *SERPINA1* and *TTR,* among others (Figure 1C, and Supp. Table 2). Differential expression analysis was used to annotate the four subgroups of hepatocytes present in all four donors and treatment conditions (Figure 1B, Supp. Table 2, Methods). Representative marker genes illustrate functional specialization. For instance, *ATF6*, *CYP8B1* and *HMGCS2* for sterol and bile acid synthesis were up-regulated in subgroup I. Subgroup II was represented by *LDHA*, *GSTZ1* and *GSTO1*, involved in the carbohydrate and phase II metabolism. Subgroup III was characterized by *ABCC2, ABCC3* and *APOB* over-expression, among other genes (Supp. Figure 2B) involved in lipid and phase III metabolism (Figure 1C). Lastly, subgroup IV was characterized by the loss of gene expression, which was 11-fold reduced compared to subgroup I (Figure 1D, Methods). The instability of PHHs has been previously characterized for the loss of expression of liver-specific transcription factors and downstream target genes [55, 60–62].

Across all conditions, 38.2% of the cells were specialized in the metabolism of bile acids and sterols (subgroup I) [59]; 38.7% of the cells were enriched in genes involved in carbohydrate metabolism and phase II enzymes (subgroup II); 5.4% of the cells were responsible for lipid metabolism and expression of phase III enzymes (subgroup III) (Supp. Figure. 2C); and 17.7% of the cells were losing expression (subgroup IV) (Supp. Figure. 2C).

Interestingly, in subgroup III, the majority of the cells (73.2%) were assigned to S-phase according to the cell cycle phase classification using the tool *cyclone* (Figure 1E, Supp. Figure 2A). Moreover, subgroup II showed a higher expression level of stress marker genes such as *RSP19*, *PRDX1, BAX, GSTA1, LGALS1, MTH1* and *MTHM* (Supp. Figure 2D, Supp. Table. 2). Likewise, subgroup II showed a low percentage of cells in which a gene is expressed, suggesting that these cells might be prone to lose their characteristic hepatocyte-like expression profile in culture.

To further characterize underlying upstream molecular events, we used the ChIP-X Enrichment Analysis tool, ChEA3 [63] to reconstruct the network of putative transcription factors (TFs) regulating gene expression. We focused our analysis on the untreated cells (i.e. DMSO-treated cells) across all donors, to prevent potential effects due to treatment conditions. Considering the top 500 differentially expressed genes per subgroup, key hepatic TFs, such as *HNF4A* [64, 65] and *MLXIPL* [66, 67] were found among the top 25 predicted TFs. Only in subgroup IV a decrease in the expression levels of key transcription factors was detected (Supp. Figure 2E, Supp. Table 3), while subgroups I, II and III were defined by a unique combination of master regulators (Supp. Figure 2E, Supp. Table 3).

### Primary human hepatocytes retain functional specialization *in vitro* in the absence of liver zonation

Among the four subgroups of hepatocytes identified *in vitro*, only subgroups I, II and III were considered functional and metabolically active. We then further investigated whether these three hepatocyte subgroups were also present *in vivo*.

First, we investigated nine human livers described in Aizarani *et al*. in 2019 [1]. Hepatocytes were extracted from the data set based on the expression of mature hepatocyte maker genes *ALB*, *TTR*, and *HNF4A*. After performing *Louvain* clustering on the *in vivo* hepatocytes (resolution 0.2), we used the transcriptional profiles of our subgroups (I, II and III) to identify hepatocyte specialization per cluster. We observed that the transcriptional profiles defining our subgroups are similarly expressed in hepatocytes *in vivo* with a correlation up to 0.94 for the top ten DEGs per subgroup (Figure 1F and Supp. Figure 3).

*In vivo*, expression profiles of hepatocytes are affected by their position in the liver lobule [1, 48, 49, 68]. Therefore, we investigated whether the three hepatocyte subgroups identified as subgroups I, II and III *in vivo*were related to liver zonation patterns (Supp. Figure 3 A). Based on their expression profile, cells were scored for zonation marker genes (Supp. Figure 3B, Methods) and assigned to three zones: pericentral, mid-zone, and periportal (Supp. Figure 3A and 3B). In subgroup I, 64.6% of cells were assigned to periportal area, 6.6% to midzone, and 46.5% to the pericentral area. The majority of cells in subgroup II, 52.3%, were assigned to the midzonal area, with 12.1% of cells in the pericentral area and 27.9% in the periportal area. Subgroup III showed an enrichment in mid- and pericentral expression profiles, 41.1% and 41.4% respectively, and 7.5% in the periportal area (Supp. Figure 3B and 3C, Methods).

Additionally, hepatocyte subgroups and zonation were found to be intertwined *in vivo* (Supp. Fig. 3D, left). For instance, *CYP27A1* in subgroup I was pericentrally enriched [48, 69, 70]; while HS*D11B1,* involved in bile synthesis [71] was distributed periportally (Supp. Table 1; Supp. Figure 3, left). Remarkably, *in vitro*, PHH retained specialization into the three functional subgroups (I, II, and III) in the absence of zonation (Supp. Table 1; Supp. Figure 3, right). In subgroup I, the CY*P27A1* and *HSD11B1* had similar expression levels in all the cells. Similarly, zonation marker genes in subgroup II and III were evenly expressed in all cells studied.

These findings were extended to additional five human donors [5] (Supp Figure 3E and 3F, Methods), showing that PHH *in vitro* retained functional specialisation in the absence of liver zonation.

### Phenotyping cocktail used to assess induction of cytochrome P450 shows differential metabolic profiles among hepatocyte subgroups

In order to study the impact of cellular heterogeneity on liver function, we further characterized the hepatocyte subgroups by assessing their response to a phenotyping cocktail. The Sanofi-Aventis five-drug cocktail was used to simultaneously monitor the expression levels of the main five cytochrome P450 genes as a readout of the metabolic capacity of primary human hepatocytes [24, 72]. Therefore, this cocktail, consisting of a mixture of individual selective substrates of CYP2D6 (metoprolol), CYP2C19 (omeprazole), CYP2C9 (S-warfarin), CYP3A (midazolam) and CYP1A2 (caffeine), was used to monitor CYP induction [73], and incubated for 66h in primary human hepatocytes (Methods).

Incubation with the five-drug cocktail did not lead to substantial differences in the number of captured cells in each condition per subgroup (Supp. Figure 4A). Upon incubation, the induction of the mRNA of the five CYPs involved in the metabolism of the five-drug cocktail was monitored showing their up-regulation in pseudobulk (Figure 2A left, Supp. Figure 4B). However, differences in the up-regulation levels were observed between subgroups (Figure 2A, right). For instance, in pseudobulk, *CYP2C9* was up-regulated 1.7-fold in Cocktail, while in subgroup III only a 1.1-fold change was detected. Likewise, *CYP3A4* was not significantly up-regulated in subgroup III, but a 4-fold increase was detected in subgroup I (Figure 2A, right).

**Figure 2:**
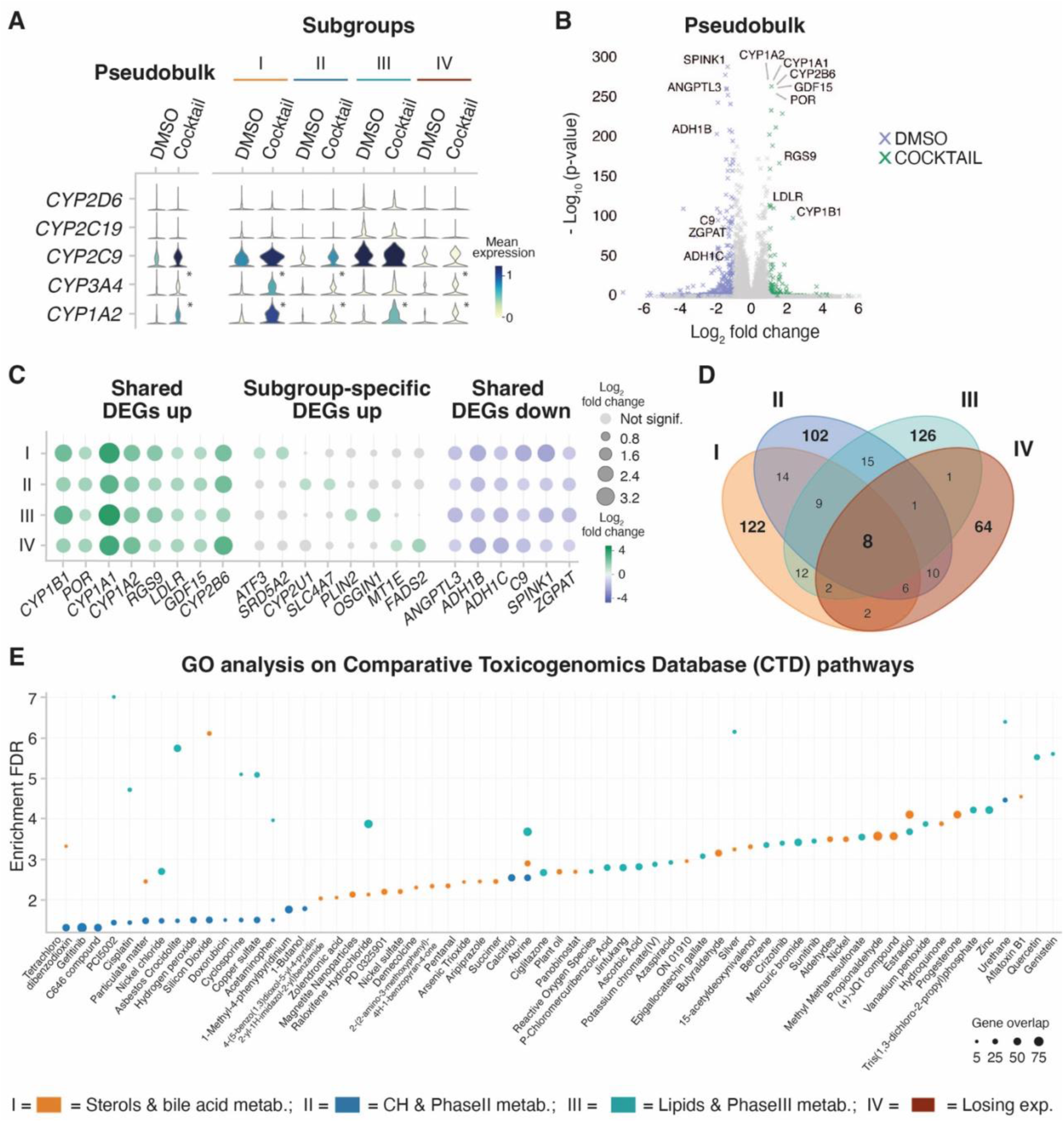
Subgroups of PHHs show different metabolic profile in response to 5-drug cocktail. **(A)** Violin plots depicting expression levels between Cocktail and DMSO of the five CYPs involved in the metabolism of the five-drug cocktail in pseudobulk (left), and in each hepatocyte subgroup (right) (*=p-value<0.05 and |log2-fold change|>1, t-test). **(B)** Volcano plot depicting the differential expression between Cocktail (green) and DMSO (purple) in pseudobulk. Text highlights the genes identified as DEGs in all subgroups. **(C)** Dot plot showing log2-fold change (color scale) and p-value (dot size) between Cocktail and DMSO for genes significantly up-regulated in all subgroups (left); representative subgroup-specific up-regulated genes (middle); and genes significantly down-regulated in all subgroups (right). **(D)** Venn diagram showing overlaps of significantly up-regulated genes upon cocktail treatment between the subgroups. **(E)** Scatter plot depicting enrichment of the genes specifically up-regulated in the metabolically active hepatocyte subgroups I, II and III in pathways known to be involved in the metabolism of given chemical compounds (Drug.CTD database). The size of the dot corresponds to the number of overlapping genes in a given pathway.

To monitor cocktail-induced changes on the transcriptome, we first analysed global changes upon five-drug cocktail incubation, detecting 161 significantly up-regulated genes compared to DMSO (Figure 2B and Supp. Figure 4C). All genes with a corrected p-value below 0.05 and a log_2_ fold change greater than 1 were designated as significantly up-regulated in cocktail-treated cells. Genes with log_2_ fold change below -1 were considered significantly up-regulated in DMSO-treated cells (Figure 2B and 2C, Supp. Figure 4D, Supp. Table 5, Methods). Additionally, removing hepatocyte subgroup IV (characterized by the loss of expression) lead to the detection of 205 significantly up-regulated genes (Supp. Figure 4C, Supp. Table 4).

Investigating the shared and subgroup-specific signatures in response to five-drug cocktail, only eight genes were significantly up-regulated upon cocktail treatment in all four subgroups: *CYP1B1, POR*, *CYP1A1*, *CYP1A2*, *RGS9*, *GDF15, CYP2A7*, and *CYP2B6* (Figure 2B and Figure 2C). These genes were also detected as significantly up-regulated in the pseudobulk analysis, and correspond to genes involved in the phase I metabolism of xenobiotics [17, 19].

Subgroup-specific DEGs differentially up-regulated were also identified (Figure 2C middle). These genes were not detected as significantly up-regulated in the pseudobulk analysis. For example, upon five-drug cocktail incubation, hepatocytes in subgroup I (bile acid and sterol metabolism) specifically up-regulated *ATF3* and *SRD5A2*; subgroup II (carbohydrate and phase II metabolism) specifically up-regulated *CYP2U1* and *SLC4A7*; and subgroup III (lipids and phase III metabolism) specifically up-regulated *PLIN2* and *OSGIN* (Figure 2C middle). Similar number of genes were specifically up-regulated in every metabolically active hepatocyte subgroup: 122 genes in subgroup I, 102 genes in subgroup II, and 126 genes in subgroup III. Meanwhile, only 64 genes were specifically up-regulated in subgroup IV (Figure 2D).

Additionally, Gene Ontology (GO) analysis of subgroup-specific DEGs was performed using the Comparative Toxicogenomics Database (CTD) to explore toxicological interactions. This database is particularly suited for drug-disease or drug-phenotype interactions [73–75]. GO analysis using CTD showed that each hepatocyte subgroup specialized in the metabolism of certain xenobiotics, based on their differential transcriptomic profile (Figure 2E, Supp. Figure 4E). For instance, the metabolic pathway of abrine was up-regulated across all hepatocyte subgroups I, II and III. However, some compounds were only enriched in two subgroups, like cisplatin, in subgroups I and II. Finally, the pathways for the metabolism of other compounds were only enriched in one subgroup, like of ciglitazone and aflatoxin B1, enriched in subgroup II and I, respectively.

In summary, these results indicate that upon five-drug cocktail treatment, hepatocytes subgroups showed differential transcriptional responses, characterized by shared metabolic pathways and subgroup-specific transcriptional profiles associated with different potential for metabolizing endobiotic (endogenous) and xenobiotic (exogenous) chemical compounds.

### Intracellular lipid accumulation leads to differential transcriptional variability among hepatocytes subgroups

Recently, it has been shown that changes in cytochrome P450 activity correlates with altered lipid metabolism [76–78]. For example, hepatic steatosis affects the transcriptomic profile of parenchymal and non-parenchymal cells as well as the cellular composition in the liver [30, 31, 79, 80]. Particularly in hepatocytes, the lipid metabolism is disrupted upon fat accumulation through an alteration of key enzymes in the lipid synthesis, storage and clearance pathways [30, 81][82]. Furthermore, an increase in the production of chemokines has been observed, associated to the inflammation occurring in NAFLD [83, 84].

To shed light on the effect of lipid accumulation on the metabolic capacity of functional subgroups of PHH, hepatic steatosis *in vitro* was modelled by incubating the cells with 200 µM mixture of a 2:1 ratio of oleic acid to palmitic acid [37] (Methods). This mixture has previously been shown to mimic benign chronic steatosis with minor lipotoxic and apoptotic effects [36–38]. First, we investigated whether - and to what extent - fat accumulation triggers a coordinated transcriptional response in PHHs. Thus, the coefficient of variation for DMSO- and FFA-treated cells was calculated per subgroup. Overall, cells losing expression (subgroup IV) showed the highest transcriptional variability (Figure 3A, Supp. Figure 5A). Moreover, lipid accumulation significantly increased the variability in functional subgroups I and II, but decreased it in subgroup III (Figure 3A). This indicated that subgroup III showed a more coordinated response towards accumulation of lipids.

**Figure 3:**
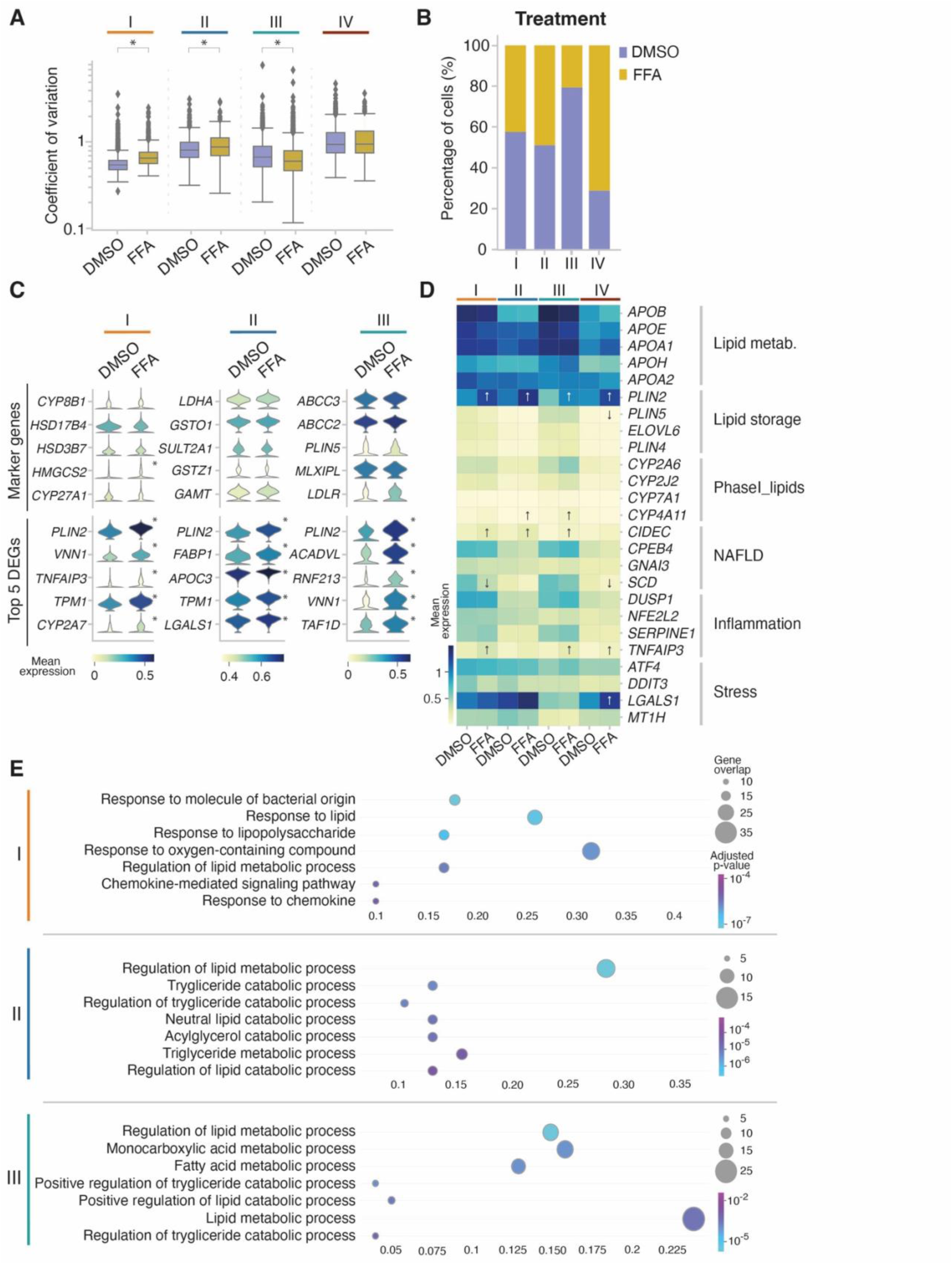
Intracellular lipid accumulation increases loss of expression and transcriptional variability. **(A)** Box plots depicting the coefficient of variation per gene in cells treated with DMSO or with FFA per subgroup (*=p-value<0.05, *MannWhitneyU*). **(B)** Bar plot showing the percentage of cells treated with DMSO or with FFA in every subgroup. **(C)** Stacked violin plots of the genes used in figure 1 to identify the subgroups, split into DMSO- and FFA-treated cells (top) and top 5 up-regulated genes upon fat accumulation per subgroup (bottom) in DMSO and FFA treatments (*=p-value<0.05 and |log2-fold change|>0.75, t-test). **(D)** Heatmap depicting the logarithmic mean expression of genes related to lipid metabolism, lipid storage, NAFLD-related genes, inflammation and stress response in DMSO- and FFA-treated cells per subgroup (↑: indicates up-regulation towards DMSO; ↓: indicates down-regulation towards DMSO, t-test). **(E)** Scatter plot of the gene ontology (GO) analyses using *gprofiler* of the genes up-regulated upon fat accumulation in each of the subgroups. The top 7 most significantly enriched terms are depicted.

Secondly, we observed that the proportion of cells treated with FFA differed between the functional hepatocyte subgroups. More than 74% of the cells in subgroup IV were FFA-treated cells, while in subgroup III 24% of the cells were FFA-treated cells. Similar percentage of FFA-treated cells were found in subgroup I and II (Figure 3B, Supp. Figure 5B).

To explore global changes triggered by lipid accumulation, differential expression analysis was performed between FFA- and DMSO-treated cells. Compared to Cocktail treated cells, fewer up-regulated genes (3.5 times) were detected in FFA-treated cells over DMSO expression level (Supp. Figure 5C). Thus, genes with a corrected p-value below 0.05 and a log_2_ fold change greater than 0.75 were designated as significantly up-regulated in FFA-treated cells. Genes with log_2_ fold change below -0.75 were considered significantly up-regulated in DMSO-treated cells (Supp. Figure 5C, Methods).

Thirdly, to track down the impact of intracellular fat accumulation on cellular identity, transcriptional changes in the previously selected marker genes - defining the hepatocyte subgroups - were investigated (Figure 3C top and Figure 1F). No significant changes in mean expression were observed for most of these marker genes upon accumulation of fat. Only *HMGCS2,* a key enzyme responsible for the synthesis of ketone bodies [85, 86] was significantly up-regulated in subgroup I upon fat accumulation (Figure 3C, top).

In the metabolically active subgroups (I-III), the top five significantly up-regulated genes under fat accumulation were known markers of lipid droplet formation and lipid metabolism (Figure 3C, bottom). For instance, the lipid droplet-associated perilipin protein *PLIN2* was significantly up-regulated in all subgroups, which has previously been shown to be relevant in diet-induced NAFLD [87–89]. The inflammation marker *TNFAIP3*, also discovered to ameliorate NAFLD and be protective against its progression [90, 91] was up-regulated among the five top DEG genes in subgroup I (Figure 3C).

Subsequent analysis of fat-metabolism related pathways and genes involved in stress response [92] showed subgroup-specific signatures upon intracellular fat accumulation (Figure 3D, and Supp. Table 2 and Supp Table 4). For example, *CYP4A11,* involved in NAFLD progression by inducing ROS-related lipid peroxidation and inflammation [93–95], was significantly up-regulated in subgroups II and III; and *CIDEC*, a promotor of triglyceride accumulation [96, 97], was significantly up-regulated in subgroups I, II and III. Cells losing expression (IV), up-regulated *LGALS1* [98, 99], and *HSPB1* [100, 101] (Figure 3D and Supp. Figure 5D), which are known markers in the gene ontology pathway GO:0006950: “response to stress”.

To identify the main biological processes associated to each metabolically active subgroup upon fat accumulation, gene ontology (GO) analyses were performed using all the significantly up-regulated genes per subgroup. Subgroup I showed gene overlaps in pathways related to cellular response to lipids, together with lipopolysaccharide and chemokine metabolism (Bode et al. 2012.European Journal of Cell Biology) (Figure 3E). For instance, chemokines *CXCL8, CXCL1, CXCL10* and *CXCL11* were up-regulated in FFA condition (Supp. Table 7), suggesting that lipid accumulation could lead to increased inflammation through chemokine signaling [102–104]. Moreover, subgroup II exhibited a high overlap of genes involved in the regulation of triglyceride metabolic processes, as well as in the acylglycerol catabolic process, denoting that these cells were rather involved in clearance of neutral lipids [105, 106]. Finally, subgroup III cells showed enrichment in lipid, monocarboxylic acid and fatty acid-related metabolic processes and lower transcriptional variability, most likely due to their coordinated response to fat accumulation [9, 107–110].

Taken together, in metabolically active hepatocyte subgroups I and II, intracellular lipid accumulation increased transcriptional variability, thus affecting the fine-tuned regulation of lipid metabolism. Importantly, in subgroup III, specialized in the metabolism of lipids, transcriptional variability was reduced upon fat accumulation, suggesting a robust and tight coordinated response to chronic accumulation of lipids.

### Intracellular lipid accumulation impairs drug metabolism phase I, II and III, with concomitant up-regulation of stress-related pathways

Large inter-individual variability has been observed in the metabolism of CYP substrates *in vivo*. This variability might be caused by genetic, epigenetic, and environmental factors. For instance, the simultaneous administration of five or more drugs, known as polypharmacy, is highly common in the clinical practice [44, 111]. The co-administration of drugs increases the risk for developing hepatotoxicity and Adverse Drug Reactions (ADRs), such as Drug-Induced Liver Injury (DILI) [112–114]. Additionally, a higher incidence of DILI has been reported in patients suffering from NAFLD [45–47]. Therefore, we assessed the impact of fat accumulation in the phase I metabolism of drugs, using the previously characterized phenotyping cocktail (Sanofi-Aventis) (Figure 2). The expression of the individual CYPs targeted by the drug cocktail was analyzed, comparing Cocktail- and FFA+Cocktail-treated cells (Figure 4A). In all subgroups, fat accumulation decreased the expression levels of the five targeted cytochromes (Figure 4A). For instance, in subgroup I, II, and IV, *CYP3A4* was significantly up-regulated upon cocktail treatment, but its induction was attenuated upon chronic lipid exposure. In subgroup III, the largest magnitude change of CYP expression was observed. Notably, *CYP2D6, CYP2C19, CYP2C9* and *CYP3A4* were significantly down-regulated in comparison to baseline DMSO level upon FFA+Cocktail treatment.

**Figure 4:**
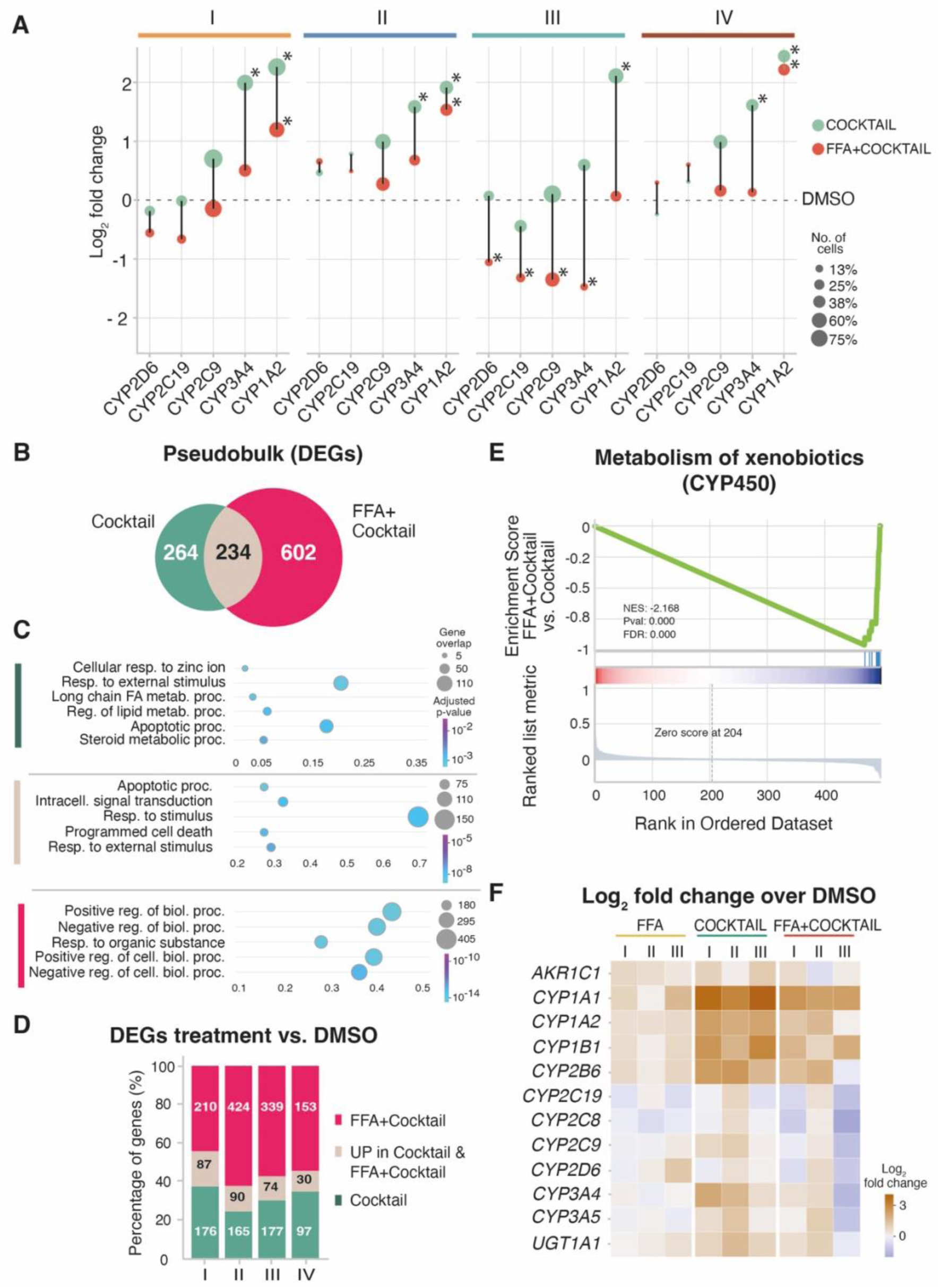
Intracellular lipid accumulation impairs drug metabolism phase I, II and III. **(A)** Scatter plot depicting the log2-fold change of the 5 induced cytochromes between cocktail and DMSO (green), and between FFA+cocktail and DMSO (red) in each subgroup (*=p-value<0.05 and |log2-fold change|>1, t-test). Dot size corresponds to number of cells in which the gene is expressed. **(B)** Venn diagram showing the overlap of DEGs between Cocktail- vs. DMSO-treated cells (green) and FFA+Cocktail vs. DMSO-treated cells (red). **(C)** Scatter plot of the gene ontology (GO) analyses using *gprofiler* of the genes up-regulated specifically upon cocktail treatment (green); up-regulated specifically upon FFA+Cocktail treatment (magenta), and in both Cocktail and FFA+Cocktail (beige). The top 5 most significantly enriched terms are depicted. **(D)** Bar plot showing for each subgroup the percentages of genes specific to: a) Cocktail vs. DMSO (green); b) Genes up-regulated in both, Cocktail vs. DMSO and FFA+Cocktail vs. DMSO (beige); c) Specific to FFA+Cocktail vs. DMSO (magenta). **(E)** GSEA plot for FFA+Cocktail vs. Cocktail on the pathway of “Metabolism of xenobiotics by CYP450”, enriched in the Cocktail vs. DMSO-specific genes. **(F)** Heatmap depicting log2-fold change to DMSO level of genes involved in drug-metabolism related pathways that were enriched in Cocktail vs. FFA+Cocktail tre

Additionally, to identify global transcriptional changes between Cocktail and FFA+Cocktail, differential expression analysis was performed in pseudobulk (Methods). Using DMSO as a baseline expression, 264 genes were up-regulated uniquely under Cocktail treatment; 234 genes were commonly up-regulated in both Cocktail treatment and FFA+Cocktail; and 602 genes were up-regulated solely in FFA+Cocktail, suggesting that more biological processes are affected (Figure 4B, Supp. Figure 4D and 6A, Supp. Tables 5 and 8). To further investigate the affected pathways, gene ontology (GO) analyses were performed on these genes (Figure 4C). Specifically upon Cocktail treatment we observed an evident enrichment in pathways responsible for the metabolism of xenobiotic compounds (Figure 4C, top, green). Genes commonly up-regulated upon Cocktail and FFA+Cocktail were less specific for drug metabolism and showed an enrichment in pathways for general stimulus responses (Figure 4C, middle, beige). Finally, the genes specifically up-regulated in FFA+Cocktail showed an enrichment in stress-related pathways (Figure 4C, bottom, magenta). The percentage of genes in each category was similar in all four subgroups of hepatocytes, suggesting that drug metabolism was similarly affected by the accumulation of fat (Figure 4D). In addition, a downregulation of key hepatic marker genes was observed in FFA+Cocktail (Supp. Figure 6B). For instance, *CYP2A7* was up-regulated in subgroups I, III and IV upon Cocktail treatment, but the induction was impaired in the presence of fat (Supp. Figure 6B).

With the aim of dissecting the impact of chronic accumulation of fat on drug metabolism, we compared FFA+Cocktail and Cocktail treated cells in a gene set enrichment analysis (GSEA)(Methods) [115, 116]. Among the top significant enriched pathways, the metabolism of xenobiotics by the CYP450 family was down-regulated (Figure 4E), while the insulin resistance pathway was up-regulated in the presence of fat (Supp. Figure 6C), indicating a dysregulation of multiple metabolic pathways. For example, while chronic exposure to lipids did not trigger a consistent up-regulation of the insulin resistance pathway, the combination of FFA+Cocktail led to the up-regulation of *NFKBIA, TRIB3, CREB3L3, IRS2, SLC2A1, SOCS3, PIK3CD, CREB5, RELA, CPT1B* in all subgroups (Supp. Figure 6D). Moreover, upon incubation with Cocktail, drug metabolism-related genes were up-regulated consistently in all functional subgroups of hepatocytes (I, II, II) (Figure 4F). However, in the FFA+Cocktail treatment, subgroup III showed a drastic down-regulation in multiple phase I CYPs, e.g. *CYP2C19, CYP2C8, CYP2C9, CYP2D6, CYP3A4, CYP3A5;* in phase II enzymes *GSTA1*, *GSTA2*, and *SULT2A1;* as well as in phase III enzymes *SLCO1B1* and *ABCG5,* exhibiting an overall impairment in all three phases of drug metabolism.

In summary, we observed that intracellular lipid accumulation led to an impairment of drug metabolism characteristic for each subgroup, in which multiple metabolic pathways are simultaneously affected. Therefore, primary human hepatocytes lose their drug-related metabolic specificity in the presence of chronic accumulation of lipids.

## DISCUSSION

The application of single-cell genomics technologies allows the dissection of subtle differences within a seemingly homogeneous population of cells, which has been demonstrated in a plethora of organs, tissues and cell types [107], including liver [1, 4, 5, 31, 49, 117–121]. The assessment of safety, toxicity and efficacy of drugs is generally performed in bulk analyses, representing average features of the most abundant transcripts or the most abundant cell type [37, 61, 62]. However, this approach hinders the assessment of cellular heterogeneity and the identification of cellular phenotypes that might be rare or display opposite transcriptional responses.

To circumvent this limitation, we have used single-cell transcriptomics to assess the metabolic profiles of individual PHHs, a gold standard human *in vitro* liver model, to study drug-related metabolic capacity in healthy condition and in response to environmental factors. Across all donors and treatment conditions, four major subgroups of hepatocytes were identified. Consistently with previous literature on PHHs [60-62, 122, 123], we identified a subgroup of PHHs losing the characteristic mature hepatocyte signature after 72h in culture (subgroup IV, Figure 1, Supp Figure 2). For instance, in this subgroup we observed a down-regulation of upstream liver enriched transcription factors such as *MLXIPL* (*ChREBP*), *RXRA*, *NH1H4* (*FXR*), *PPARA*, *HNF4A* and *CEBPA* [64, 65, 124, 125] (Supp. Figure 6B) and low expression of the transcription factors predicted by the ChEA3 tool (Supp. Figure 2E). Further analysis of metabolic profiles showed that subgroups I, II and III were specialized in sterol and bile acid, carbohydrate and phase II, and lipids and phase III metabolism, respectively. Thus, we defined subgroups I, II and III as metabolically active and functional subgroups (Figure 1). Interestingly, in subgroup II, a high expression of stress markers was observed, suggesting that this group is prone to eventually lose mature hepatocyte-like expression (Supp. Figure 2) [126–128]. This observation is supported by the high correlation between subgroup II and subgroup IV *in vitro* (Supp. Figure 3F). Moreover, subgroup I and II showed a larger percentage of cells *in vitro*, followed by subgroup IV and III (Supp. Figure 2C), implying that *in vitro* functional specialization and degree of heterogeneity depend on the proportion of hepatocytes subgroups and human variability.

Another source of functional specialization is the well-known hepatic zonation in the liver. The zonation patterns found *in vivo* due to the oxygen and nutrient gradient along the lobule axis are not conserved in 2D *in vitro* models [129–135]. To deeply investigate if hepatocyte subgroups are related to reminiscent liver zonation, we have compared our findings *in vitro* with two *in vivo* datasets comprising nine and five human donors, respectively [1, 5]. We have revealed the presence of similar functional hepatocyte subgroups *in vivo*, showing that the three functional subgroups identified are mostly independent of zonation. Complementarily, marker genes used for subgroup identification are subjected to zonated expression *in vivo* (Supp Figure 3) [1, 4, 5, 10, 48].

In order to study the impact of these four subgroups of hepatocytes in liver function, we have further characterised their drug metabolic capacity by assessing their functional response to a phenotyping cocktail (Figure 2). The drug metabolism *in vitro* can be defined by three major phases. In phase I, oxidation, hydrolysis, reduction and cyclization reactions are catabolized mainly by the cytochrome P450 (CYP450) superfamily of monooxygenase enzymes. The main members are the isoforms CYP1A2, 2C9, 2C19, 2D6 and 3A4, that account for the metabolism of around 70-80% of the clinically available drugs [17, 19, 136]. The expression of these cytochromes is induced by the presence of their substrate compounds, which are generally used as a measurement of liver metabolic capacity. These substrates act as metabolic probe to assess the functional phenotype of the liver. This strategy, formally known as the “cocktail approach”, has been used to monitor the cytochrome enzymatic activity and changes in the induction of their corresponding mRNA levels (Berger e*t al. 2*016; Fuhr e*t al. 2*007; Ryu e*t al. 2*007; Turpault e*t al. 2*009) Bosilkovska e*t al. 2*014; Chainuvati e*t al. 2*003; Christensen e*t al. 2*003). Among several drug-cocktail approaches, we have selected the Sanofi-Aventis cocktail due to its versatile usage in murine [72, 137], primates [138, 139], dogs and minipigs [138, 140], and human studies [24], both *in vivo* [24, 72, 139] and *in vitro* [138].

The Sanofi-Aventis cocktail revealed that the identified hepatocyte subgroups responded differently to the metabolic challenge at the transcriptomic level, and that these differences cannot be anticipated through analyses performed in bulk (Figure 2). For instance, a high percentage of cells in subgroup IV might lead to an underestimated effect of drug induction, which is highly relevant in the safety evaluation of xenobiotics [141]. Additionally, exploration of toxicological interactions (i.e. CTD) showed that each hepatocyte subgroup is specialized in the metabolism of certain xenobiotics, which suggests that certain subgroups of hepatocytes could be more susceptible to develop Adverse Drug Reactions (ADRs) and toxic metabolites when challenged by a specific compound (Figure 2E). Furthermore, we anticipate that chronic liver diseases might also affect the percentage of cells in each hepatocyte subgroup and therefore have major implications in the assessment of drug efficacy, safety and toxicity in the early phases of drug development.

To assess impact of chronic liver disease on cellular heterogeneity, we have mimicked hepatic steatosis and early stages of non-alcoholic fatty liver disease (NAFLD) by loading the cells with intracellular lipids (Figure 3) [37, 142, 143]. Chronic accumulation of fat led to an increase in transcriptional variability in subgroups I and II, indicating transcriptional noise and random fluctuation in the mRNA level in individual cells [144]. Importantly, in subgroup III, specialized in the metabolism of lipids, transcriptional variability was reduced upon fat accumulation, suggesting a robust and tight coordinated response to chronic accumulation of lipids. Upon chronic lipid accumulation, the number of cells losing their characteristic expression increased suggesting loss of functionality. Additionally, subgroup-specific signatures were identified upon intracellular fat accumulation, in which lipopolysaccharide and chemokine metabolism was upregulated in subgroup I (Figure 3). The upregulation of chemokines in the most abundant hepatocyte subgroup suggests that lipid accumulation could lead to increased inflammation in a specific cell population. These results reinforce the notion that changes in the proportion of hepatocyte subgroups might determine the functional outputs in response to environmental or dietary factors.

For instance, hepatic fat accumulation also occurs during heathy ageing [161], where increased transcriptional variability [9, 109] in several tissues and cell types has been observed [9, 107, 145–147] . Additionally, ageing affects the hepatocyte function [148] and cytochrome P450 enzymes [42, 149–152]. Therefore, further investigations on the age-associated cellular heterogeneity and drug metabolism could anticipate the unexpected adverse drug reactions or drug-induced liver toxicity in the elderly.

Additionally in the elderly, the co-administration of drugs known as polypharmacy is highly common for the treatment of age-related comorbidities, increasing the risk for the development of ADRs and more specifically, DILI [41–43][44]. In all subgroups of hepatocytes, intracellular lipid accumulation diminished their capacity to metabolize the five-drug cocktail (Figure 4). Especially, subgroup III, responsible for lipid and phase III metabolism, showed the most prominent decrease in the five targeted cytochromes simultaneously (Figure 4). Importantly, intracellular lipid accumulation led to an impairment of drug metabolism characteristic for each subgroup, in which multiple metabolic pathways are simultaneously affected. In summary, cellular heterogeneity is affected by intrinsic and extrinsic factors which might lead to aberrant metabolism, producing toxic metabolites and associated stress-related responses and inflammation.

On the one hand, intracellular lipid accumulation increases transcriptional variability and loss of expression *in vitro,* on the other hand it diminishes hepatic capacity for drug metabolism in individual cells. Translating these results to *in vivo* human biology, our findings indicate that fat accumulation could further accelerate age-related processes by means of increased transcriptional noise and susceptibility to adverse drug reactions. Finally, our results suggest that hepatic steatosis leads to changes in cellular composition affecting tissue function. Assessing the impact of cellular heterogeneity on tissue function will shed light on novel molecular mechanisms underlaying chronic or age-related pathologies.

## MATERIAL AND METHODS

An outline of the experimental design is shown in Figure 1. Briefly, commercially cryopreserved Primary Human Hepatocytes (PHH) were purchased from Lonza from four different donors: HUM180812 (male, 57 years old, Hispanic) and HUM4152 (male, 18 years old, Caucasian), HUM181641 (male, 56 years old, Caucasian) and HUM4190 (male, 26 years old, Caucasian). All donors had a Body Mass Index in the normal range and were not diabetic (Supp. Table 1). The first two donors were processed simultaneously in a first batch, and the second two in a second batch, aiming for a maximum of eight samples processed at a time.

### Cell culture

Each cryovial of PHH was thawed and plated according to Lonza’s “Suspension and plateable cryopreserved hepatocytes: technical information and instructions.”. the protocol was followed stepwise minutely, using the recommended thawing and plating media (Lonza, MCHT50 and MP250, respectively). The cells were dispensed and mixed using only wide orifice tips (Rainin, Ref. 17014297). For efficient cell seeding densities and attachment, cells were counted using Trypan Blue Exclusion Method and seeded into Collagen-I coated plates at a density of approximately 100,000 cells/cm^2^ following the Lonza’s instructions (Lonza, “Suspension and Plateable Cryopreserved Hepatocytes Technical Information and Instructions”). Six hours post seeding, cells were washed with 1 mL of pre-warmed Maintenance Medium (Lonza, MCHT50) before addition of treatment media. The treatment medium was renewed every 24h for a total incubation period of 72h post-seeding. Free fatty acids (FFA) consisting of a 200 µM mixture of a 2:1 ratio of the unsaturated oleic acid to the saturated palmitic acid were added to the maintenance media, to mimic the levels in human steatosis (Gómez-Lechón et al. 2007). In order to facilitate FFA uptake, pre-bounding of free fatty acids to 1% bovine serum albumin in a 1:5 molar ratio (Sigma-Aldrich) was performed by heating the mixture at 40°C for 2 hours (Kozyra *et al.* Sci Rep 2018).

### Drug cocktail preparation and storage

The individual components of the 5-drug cocktail[24] were dissolved in sterile DMSO, filtered through a 0.2 µM syringe filter (Merck, SLGVV255F) and stored at -80°C for a maximum of six months (“compound stock concentration”, Supp. Table 10). The individual drugs were mixed at 200x concentration (“working concentration”, Supp Table 10) and added to the cells to a final concentration of 80µM Caffeine (Sigma-Aldrich, Ref. 56396-100MG), 5µM Midazolam (LGC Chemicals, Ref. LGCFOR1106.00), 17µM Omeprazole (TRC Chemicals, Ref. 0635000), 20µM S-Warfarin (Sigma Aldrich, Ref. UC214-5MG) and 23µM Metoprolol (TRC Chemicals, Ref. M338815). The final DMSO concentration used on the cells was 0.5% (v/v), in all conditions.

### Single-cell RNA-seq sample preparation and sequencing

After a total of 72-hour incubation in treatment culture medium, cells were detached with pre-warmed 0,25% Trypsin (Gibco, 25200056) for 7 minutes, followed by trypsin inactivation. The dissociated cells were then collected to pellet by centrifugation at 100x *g* for 5 min at room temperature (RT). Cells were washed twice with 1 mL of pre-warmed 1xPBS pH 7.4 (Gibco, 10010023), followed by cells pelleting at 100 x *g* for 5 min at RT. A live-cell selection was performed using Dead Cell Removal Kit (Miltenyi Biotec, 130-090-101) as follows: cells were pelleted at 100x *g* for 5 min at RT, resuspended in 100 µL of dead-cell removal microbeads and incubated for 15 min at RT. Dead cells were positively selected by passing the cell suspension through a MS column and performing a wash with a total of 2 mL of Binding buffer. Living cells were eluted from the column and collected in 2 mL Eppendorf tubes. After pelleting by centrifugation at 100x *g* for 5 min at RT, cells were resuspended in 1xPBS pH 7.4 supplemented with 0.04% BSA (Miltenyi Biotech, 130-091-376), stained with trypan blue to assess viability, and counted in a hemocytometer.A single-cell suspension was obtained by dissociating cells with wide orifice pipette tips and preparing the target cell stock concentration for loading the 10X chip.

Single-cell RNA-seq libraries were prepared from each sample following the 10X Genomics Single Cell 3’ Reagent Kit User Guide (v3 or v3.1, manual CG000183 and CG000204, respectively) in the Chromium Controller (10X Genomics). The quality control of cDNA and obtained final libraries was performed using a Bioanalyzer High Sensitivity DNA Analysis assay (Agilent). Library quantification was performed using the Collibri™ Library Quantification Kit (Thermo Fischer Scientific, A38524500) in a QuantStudio™ 6 Flex Real-Time PCR System (Thermo Fisher Scientific).Both batches were sequenced in a NovaSeq6000 sequencer (Illumina) at the HMGU Core Facility for NGS Sequencing. The first batch (HUM180812 and HUM4152) was sequenced in a S2 flowcell at a depth of 250,000 reads per cell. The second batch (HUM181641 and HUM4190) was sequenced in a SP flowcell at a sequencing depth of 20,000 reads per cell. The sequencing length was set as indicated by 10X Genomics: 28/8/0/91.

### Read alignment, counting and filtering of the combined batches

Reads were aligned to GRCh38 and counted using 10X Genomics Cellranger 4.0.0 with standard parameters for each batch individually (Supp. Table 2). The resulting count matrices were combined into a count matrix of 63,527 cells times 19,971 genes. Cells with at least 1,000 counts and 500 genes were kept. Genes were kept if they were present in at least 5 cells and had fewer than 5 million reads. *Scrublet* [153] was used to identify doublets. Due to hepatocytes being subject to polyploidyzation, a lenient cutoff of 0.15 was used to avoid unwanted removal of tetraploid hepatocytes. This led to 1.7% of cells being annotated as doublets and subsequetly being removed. Lastly, cells with more than 1% mitochondrial reads were removed, resulting in a filtered matrix of 49,378 cells times 16,256 genes.

### Normalization and initial clustering

*Scran* was used for library size normalization with parameter *min.mean=0.05*. After normalization, cells with more than 20,000 normalized counts were removed. *Scanpy* functions were used for calculating principal components (PCs) and clustering. Shortly, the top 50 PCs were used to construct a neighboring graph before calculating a UMAP embedding and *Louvain* clusters. For two samples, encapsulation of the cells in droplets partially failed during the 10X library preparation (wetting failure). Therefore, an initial *Louvain* resolution of 0.5 was used to verify that cells stemming from these samples clustered separately. Indeed, some cells from one of the failed samples clustered apart from the rest of cells, and another two clusters showed fractal structures that were not present in the samples where encapsulation worked correctly. Hence, based on the knowledge of the wetting failure, these three clusters were removed, resulting in a final matrix of 38,232 cells times 16,256 genes. To remove batch effects in downstream analysis, the two batches were integrated using *Harmony* [154].

### Subgroup annotation based on individual donor analysis and clustering

In each of the four donors, the resolution of the *Louvain* clustering was selected so that every cluster contained every treatment condition (Supp. Figure 2A). Then, the top 1,000 genes per *Louvain* cluster were calculated using *sc.tl.rank_genes_groups* with *n_genes=1000*. To find out which clusters were shared between donors, the percentage of overlaps between the top 1,000 genes per cluster was calculated and hierarchical clustering was performed. By this approach, three distinct groups of similar *Louvain* clusters were detected between donors (Supp. Figure 1C). Two of the donor-specific clusters did not group together with clusters from other donors. Therefore, they were labeled as individual clusters to inspect where they would fall on the integrated data set (Supp. Figure 1C). Due to differences in sequencing depth, one group was identified in the first batch which was assigned to the cells losing expression. After integration and putting the group labels from the individual donor analysis on top of the combined UMAP, *Louvain* clustering was performed, showing that cells from the second batch that we had assigned to subgroup II were clustering with the cells losing their expression from the first batch. Therefore, those cells we re-labelled as losing their expression, subgroup IV (Supp. Figure 2A). One of the two donor-specific clusters that could initially not be assigned to a shared cluster was split into two clusters on the combined data set, which were assigned to subgroup I and subgroup II, respectively. The other individual cluster could be associated to the cells losing their expression. Marker genes were taken from reported literature to assign metabolic preferences to the functional subgroups I, II, and III (Supp. Table 2).

### Comparison to *in vivo* data

Data were obtained from GEO (Accession number GSE124395) [1]. Genes with zero expression across all cells, and cells that did not express any genes, were removed. During further filtering, cells with 100 to 6,000 genes and 800 to 30,000 reads were kept and genes were kept if they were covered in at least 10 cells, resulting in a count matrix of 11,059 cells times 19,416 genes. To keep the processing comparible to our data, normalization was performed using *scran* with parameter *min.mean=0.05*. After normalization, cells with more than 20,000 normalized counts were further removed, resulting in a count matrix of 11,043 cells times 19,416 genes. Clustering was performed using *scanpy* functions as described above. *Louvain* clustering was performed at resolution of 0.08 to computationally separate the cell types in this data set from each other. Expression of mature hepatocyte markers, such as *ALB, HNF4A* and *TTR* was used to identify the hepatocyte cluster. *Louvain* clustering with resolution of 0.2 was then performed on the hepatocytes to identify hepatocytes subgroups *in vivo*. The expression of marker genes used for subgroup identification in our data was investigated on the *Louvain* clusters. Based on this marker gene expression, *Louvain* clusters were assigned to subgroups I, II and III (Figure 1D, Supp. Figure 3A).

As zonation impacts gene expression along the pericentral-periportal axis *in vivo*, we investigated its connection to our subgroups. Zonation markers were taken from Aizarani *et al.* 2019. Based on the 35 zones reported in their study, the genes were grouped into three zones, pericentral, mid, and periportal through binning. Then *sc.tl.score_genes* was used to calculate scores for each of the three zones (Supp. Fig. 3C). Cells were assigned to pericentral (Thummel *et al.* J Pharmacol Exp Ther 1994) if they had a CV score > 0.45 and a periportal (PV) score < 1. If cells had a PV score >= 0.65 and a CV score <= 0.45, they were assigned to periportal. The rest of the cells were assigned to mid-zone (Supp. Figure 3B and C). For each of the three subgroups, the percentage of cells in CV, mid, and PV was calculated subsequently (Supp. Fig. 3B). For visualization purposes, the subgroups were separated and zonation marker genes for each subgroup were depicted on a UMAP for the *in vivo* data set and for our *in vitro* data (Supp. Figure 3D).

To further investigate the presence of our four hepatocyte subgroups *in vivo*, an additional human data set was downloaded from GEO (Accession number GSE115469) [5] filtered and normalized as before. In order to reduce potential noise, genes were removed if they were not present in at least three cells and counts were log-transformed for better comparability between datasets. Hepatocytes were isolated based on the authors’ annotation available at GSE115469. S*canpy* functions were used as described above to perform clustering with *Louvain* resolution of 0.5 to separate subgroups of hepatocytes, leading to six *Louvain* clusters highly overlapping with the clusters reported in the initial study. Marker gene expression of the three identified subgroups in our data was again used to assign the six *Louvain* clusters to the three subgroups. To increase power, the two *in vivo* data sets were integrated using *scGen* [155] (Supp. Figure 3E). The top 10 DEGs per our *in vitro* subgroups were calculated and their scaled mean expression was correlated to the combined *in vivo* scaled mean expression. This showed high correlation of marker genes between identified subgroups *in vitro* and *in vivo* (Supp. Figure 3F).

### Differential expression analysis between conditions in the subgroups

After defining metabolically functional subgroups in our *in vitro* data, we first calculated the top 500 DEGs per subgroup under DMSO in comparison to each other. To assess what drives the basal functional specialization of the subgroups, these top 500 genes per subgroup were taken as input for the online tool ChEA3 [63] which predict transcription factors regulating the gene expression per subgroup. For the purpose of visualization, five transcription factors among the top 25 per subgroup were depicted as a stacked violin plot (Supp. Figure 2E). To explore the implications of the functional specialization on metabolic capacity of hepatocytes, we then performed differential expression analysis between Cocktail- and DMSO-treated cells in each of the functional subgroups. Genes were defined significantly up-regulated if they had a log_2_-fold change of greater than 1 and a Bonferroni-adjusted p-value below 0.05. We used the venn package to visualize overlap of significantly up-regulated genes between the subgroups. Further, ShinyGO [156] was used to investigate enrichment of the subgroup-specific up-regulated genes in pathways known to be relevant for drug metabolism (Supp. Table 4).

When comparing FFA- with DMSO-treated cells, we observed that there were on average 3.5- times fewer genes with a positive log_2_-fold change than when we compared Cocktail and DMSO. Therefore, to capture the subtler effects of fat accumulation on the cells, a gene was identified as significantly up-regulated if it had a Bonferroni-adjusted p-value below 0.05 and a log2-fold change greater than 0.75. We used the same marker genes as described above (Supp. Table 2) to count overlaps between groups of marker genes and significantly up-regulated genes in the subgroups. To investigate in which pathways related to biological processed the genes up-regulated upon FFA-treatment were enriched in every subgroup, gprofiler was used (https://pypi.org/project/gprofiler-official/).

### Assessing transcriptional variability through the coefficient of variation

It is generally assumed that lowly expressed genes have an inflated transcriptional variability. Therefore, genes with a mean normalized log-transformed expression smaller than 0.25 were removed and the coefficient of variation [157] per condition in each of the subgroups was calculated on 3,434 genes. To obtain the coefficient of variation on normalized, log-transformed counts, we used the formula described in Canchola *et al*. [158].

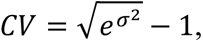

where σ² is the variation of gene *j* in the group of interest.

In every subgroup, a MannWhitneyU test was performed to check if the coefficient of variation differed significantly between treatment condition [159].

## ABBREVIATIONS

ADR: Adverse Drug Reactions
BMI: Body Mass Index
ChEA3: ChIP-X Enrichment Analysis 3
CV: Central vein
CYPs: Cytochrome P450 genes
DEGs: Differentially Expressed Genes
DILI: Drug Induced Liver Injury
FFA: Free Fatty Acids
GO: Gene Ontology
GSEA: Gene Set Enrichment Analysis
NAFLD: Non-Alcoholic Fatty Liver Disease
ns: not significant
PCA: Principal Component Analysis
PCs: Principal Components
PHHs: Primary Human Hepatocytes
PV: Portal vein
scRNA-seq: Single-cell ribonucleic acid sequencing
Supp.: Supplementary
t-SNE: T-distributed stochastic neighbor embedding
TF: Transcription Factor
UMAP: Uniform Manifold Approximation and Projection for Dimension Reduction
2D: Two-dimensional
3D: Three-dimensional

## SUPPLEMENTARY INFORMATION

Supp. Figure 1

Supp. Figure 2

Supp. Figure 3

Supp. Figure 4

Supp. Figure 5

Supp. Figure 6

Supp. Table 1 Metadata donors

Supp. Table 2 Marker genes

Supp. Table 3 ChEA3 results per subgroup

Supp. Table 4 Unique DEGs per subgroup upon Cocktail

Supp. Table 5 Differential expression Cocktail vs. DMSO

Supp. Table 6 Significance levels coefficient of variation

Supp. Table 7 Differential expression FFA vs. DMSO

Supp. Table 8 Differential expression FFA+Cocktail vs. DMSO

Supp. Table 9 Genes up-regulated upon Cocktail with and without FFA

Supp. Table 10 Drug Cocktail preparation

## DATA AVAILABILITY

All raw sequencing data is deposited in ArrayExpress [160] under accession numbers E-MTAB-11530. Additional publicly available data sets used for subgroup identification *in vivo* were obtained from GEO (Accession numbers GSE124395 [1] and GSE115469 [5])

## CODE AVAILABILITY

The jupyter notebooks containing the code to reproduce the analysis results are publicly available at https://github.com/celiamtnez/precision_toxicology. Python libraries used for the analysis include scanpy (v1.7.2), anndata (v0.7.6), matplotlib (v3.4.1), pandas (v1.2.4), numpy (v1.19.2), seaborn (v0.11.1), and scipy (v1.6.2).

## ACKNOWLEDGMENTS

This research was supported by the Helmholtz Pioneer Campus (E.S-Q, M.L.R., C.P.M-J.,), Impuls- und Vernetzungsfonds of the Helmholtz-Gemeinschaft (VH108 NG-1219 to M.C-T). We thank the Core Facility Genomics at HMGU for sequencing (I. de la Rosa) and bioinformatics (T. Walzthoeni) support, in particular Xavier Pastor for initial data analysis. We are grateful to Dr. M. Hartman, Dr. A. Schröder and Ms. A. Barden (Helmholtz Pioneer Campus) for their legal, managerial and administrative support.

## AUTHORS CONTRIBUTIONS

E.S-Q and C.P.M-J, designed experiments; E.S-Q performed experimental analyses; M.L.R. performed computational analyses. E.S-Q, M.L.R. M.C-T and C.P.M-J interpreted the data. E.S-Q, M.L.R., M.C-T and C.P.M-J wrote the manuscript. E.S-Q and C.P.M-J provided figure editing and design. All authors commented on and approved the manuscript.

